# NRF2 deficiency promotes ferroptosis of astrocytes mediated by oxidative stress in Alzheimer’s disease

**DOI:** 10.1101/2023.03.12.532248

**Authors:** Zhi Tang, Zhuyi Chen, Guo Min, Yaqian Peng, Yan Xiao, ZhiZhong Guan, Ruiqing Ni, Xiaolan Qi

## Abstract

Oxidative stress is involved in the pathogenesis of Alzheimer’s disease (AD), which is linked to reactive oxygen species (ROS), lipid peroxidation, and neurotoxicity. Emerging evidence suggests a role of nuclear factor (erythroid-derived 2)-like 2 (Nrf2), a major source of antioxidant response elements in AD. The molecular mechanism of oxidative stress and ferroptosis in astrocytes in AD is not yet fully understood. Here, we aim to investigate the mechanism by which Nrf2 regulates the ferroptosis of astrocytes in AD. Postmortem frontal cortex tissues from patients with AD and nondemented controls and brain tissue from the 3×Tg AD mouse model and wild-type mice (10 months old) were used. Immunofluorescence staining for Nrf2, the ROS marker NADPH oxidase 4 (NOX4), and GFAP was performed. We further induced Nrf2 deficiency in mouse astrocytes by using RNAi and assessed the changes in ROS, ferroptosis, lipid peroxidation, and mitochondrial dysfunction by using western blotting and immunofluorescence staining. We found decreased expression of Nrf2 and upregulated expression of NOX4 in the frontal cortex from patients with AD and in the cortex of 3×Tg mice compared to control mice. We demonstrated that Nrf2 deficiency led to ferroptosis-dependent oxidative stress-induced ROS with downregulated heme oxygenase-1 and glutathione peroxidase 4 and upregulated cystine glutamate expression. Moreover, Nrf2 deficiency increased lipid peroxidation, DNA oxidation, and mitochondrial fragmentation in mouse astrocytes. In conclusion, these results suggest that Nrf2 promotes ferroptosis of astrocytes involving oxidative stress in AD.

## Introduction

Alzheimer’s disease (AD) is the most common type of dementia, featuring deposition of amyloid-beta (Aβ) plaques, hyperphosphorylation of tau, gliosis, and progressive loss of neurons and synapses [1, 2]. Astrocytes are the most abundant glial cells in the central nervous system (CNS) and support brain physiological functions such as synapse formation and function, neurotransmitter uptake and release, and neuronal survival [3, 4]. Reactive astrocytes show disease-associated features in the brains of patients with AD and disease animal models, such as morphological and associated metabolic, antioxidant, and synaptic supporting functions [5-8]. Oxidative stress damage plays a role in the pathogenetic cascade of events in AD, even before the accumulation of Aβ or tau deposits [9-13]. The high oxygen consumption rate and abundant lipid content in the brain, compared with other organs, renders the brain particularly vulnerable to free radicals [14]. Decreased levels of expression and activities of antioxidant enzymes and an increased number of oxidative stress markers have been shown in AD brains [15, 16]. There is a vicious cycle between oxidative stress and Aβ: Aβ accumulation leads to oxidative damage, whereas pro-oxidants promote Aβ production [17, 18].

Ferroptosis is an iron-dependent programmed cell death implicated in the pathogenesis of AD [19]. Ferroptosis can be driven by uncontrolled lipid peroxidation [20, 21] and the imbalance between oxidative stress and antioxidant systems [22]. The ferroptosis pathway can be activated by intracellular inhibition of the heme oxygenase-1 (HO-1) and glutathione peroxidase 4 (GPX4) pathways and extracellular inhibition of the cystine/glutamate antiporter xCT [23]. The role of mitochondria in regulating ferroptosis has not been fully elucidated. Mitochondrial dysfunction and free radicals generated as by-products of the mitochondrial electron transport chain lead to iron dyshomeostasis and lipid peroxidation, while mitochondria-localized defense systems can also protect against ferroptosis [24, 25].

Nuclear factor erythroid 2-related factor 2 (Nrf2) is a key regulatory transcription factor that mediates the expression of a vast number of genes by binding cellular antioxidant elements in both neurons and astrocytes [26]. Under physiological conditions, Nrf2 exists in the cytoplasm as an inactive complex. When cells are stimulated, Nrf2 can translocate to the nucleus, where several endogenous antioxidant enzymes, such as nicotinamide adenine dinucleotide phosphate (NADPH) and heme oxygenase-1 (HO-1), can be further regulated by Nrf2 in the cascade reactions of oxidative stress damage [27]. Consequently, these antioxidant actions could improve mitochondrial function and prevent oxidative damage. Nrf2 has thus emerged as a key contributor to AD pathology and a therapeutic target for AD [28-30]. Nrf2 overexpression protected against neurotoxicity of Aβ and oxidative stress damage in neuronal cell culture [31-33]. In astrocyte culture, the NRF2/antioxidant responsive element (ARE) pathway negatively regulates beta-secretase 1 (BACE1) expression [34].

Here, we hypothesize that atroglial Nrf2 reduction leads to increased ferroptosis, ROS production and lipid peroxidation in AD. We evaluated the alterations in astroglial Nfr2 levels in postmortem frontal cortex tissue from patients with AD and in cortex tissue from 3×Tg mice. We assessed whether Nrf2 silencing can modulate ROS, ferroptosis, mitochondrial fragmentation and redox homeostasis in mouse astrocytes.

## Materials and methods

### Reagents, antibodies and cell cultures

Detailed information on the antibodies and reagents used in the present study are listed in **STable 1** and **STable 2**. The mouse astrocytes (mAS, M1800-57) used in this experiment were purchased from ScienCell Research Laboratories (Carlsbad, USA) and were cultured in Dulbecco’s modified Eagle’s medium (DMEM) supplemented with 10% fetal bovine serum and 100 U/mL penicillin/streptomycin at 37 □ in a humidified atmosphere of 5% CO_2_.

### Lentivirus construction and infection

Short-hairpin RNA (shRNA) targeting the Nrf2 gene (sequence: 5’-gcCTTACTCTCCCAGTGAATA-3′) was designed. Oligonucleotides were synthesized by GeneChem (Shanghai, China). Then, oligonucleotides were annealed and inserted into the pGV493-CMV-RNAi lentiviral vector (GeneChem). In brief, mAS cells were grown to 50–70% confluence in 6-well plates, and then lentivirus expressing Nrf2-shRNA was added to the target plate (MOI = 100, and 10 µl lentivirus was used per well). After culturing for another 3 days, 4 μg/ml puromycin was used to screen mAS cells transfected with stable lentivirus for 2 weeks.

### Postmortem human brain tissue

Six AD cases each with a clinical diagnosis confirmed by pathological examination of amyloid and tau (mean age 81.5, Braak stage 4-6, amyloid B-C) and six nondemented controls (mean age 76, Braak 1-2, amyloid O-B) were included in this study (detailed information in **Table 1**). Paraffin-embedded autopsy frontal cortex tissues were obtained from the Netherlands Brain Bank (NBB), Netherlands. All materials had been collected from donors or from whom written informed consent for a brain autopsy and the use of the materials and clinical information for research purposes had been obtained by the NBB. The study was conducted according to the principles of the Declaration of Helsinki and subsequent revisions. All experiments on autopsied human brain tissue were carried out in accordance with ethical permission obtained from the regional human ethics committee in Guiyang Hospital and the medical ethics committee of the VU Medical Center for NBB tissue.

**Table 1.**
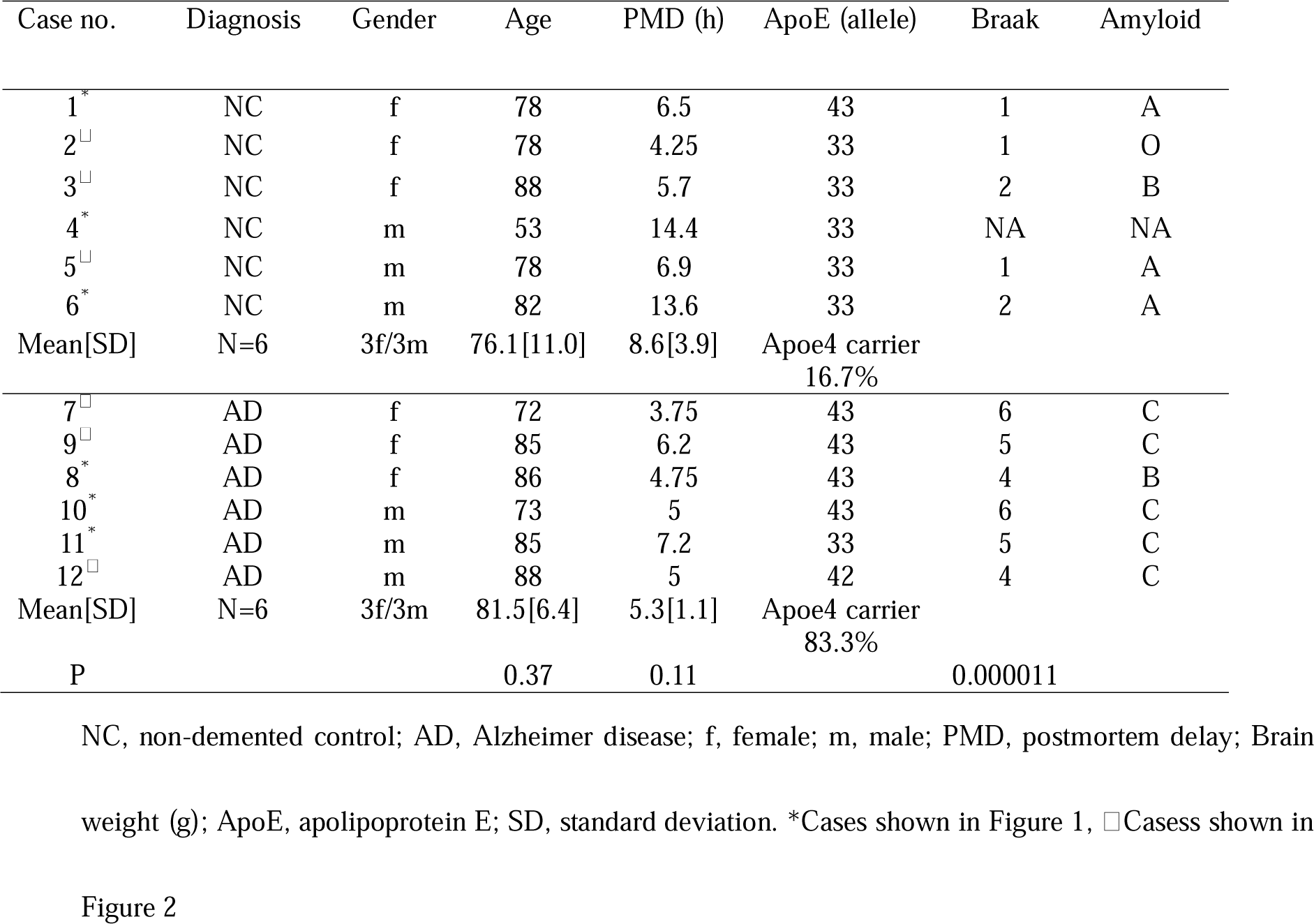
Demographics of nondemented control and AD brain tissues used in this study.

### Animal models

Five 10-month-old male transgenic APP/PS1/tau triple-transgenic mice (Stock No. 004807, B6; 129-Tg (APPSwe, tauP301L) 1Lfa *Psen1^tm1Mpm^*/Mmjax) from Jackson Laboratory (Bar Harbor, ME USA) were used [35]. Five male wild-type mice (#101045, B6129SF2/J) (male, weight 20–30 g, 10 months of age) were purchased from Jackson Laboratory as controls (Bar Harbor, ME USA). Animals were housed in individually ventilated cages inside a temperature-controlled room under a 12-hour dark/light cycle with ad libitum access to food and water. All experiments were performed in accordance with guidelines under the approval of the Animal Protection and Use Committee of Guizhou Medical University. All mice were deeply anaesthetized with pentobarbital sodium (50 mg/kg body weight, *i.p.* bolus injection) and transcardially perfused with ice-cold 0.1 M phosphate-buffered saline (PBS, pH 7.4). After perfusion, the mice were decapitated, and the brains were quickly removed. The right mouse brain hemisphere was fixed in 4% paraformaldehyde in 1× PBS (pH 7.4) for 24 h and stored in 1× PBS (pH 7.4) at 4°C. The fixed right brain hemisphere tissues were dehydrated using a vacuum infiltration processor (Excelsior ES, Thermo, USA) and embedded in paraffin using an Arcadia H heated embedding workstation (Leica, Germany).

### Immunofluorescence and confocal imaging

Paraffin-embedded frontal cortex tissue blocks from nondemented controls and patients with AD were cut at 6 µm using a microtome (HM340E, Thermo Fisher Scientific, USA). Coronal sections of the mouse brains were cut at 6 μm using a microtome. The tissue samples were deparaffinized and rehydrated prior to an antigen retrieval step (citrate buffer pH 6.0) in a microwave for 20 min at 98°C [36, 37]. The cortical sections were blocked in Tris-buffered saline with Tween-20 (TBST) with 5% normal goat serum for 30 min. Sections were then incubated with primary antibodies, including anti-GFAP, anti-NOX4, and anti-Nrf2 (**STable. 1**) at 4°C overnight. After washing in Tris-buffered saline (TBS), immunoreactions were detected using AlexaFluor488 (1:500, Life Technology, Carlsbad, CA, USA)-orAlexaFluor546 (1:500, Life Technology, Carlsbad, CA, USA)-conjugated secondary antibodies. Sections were mounted with vector anti-fading mounting medium with 4′,6-diamidino-2-phenylindole (DAPI) (Vector Laboratories, Burlingame, CA, USA). Images of human brain tissue sections were obtained by using a confocal microscope (Olympus, Japan) at 100× magnification in randomly selected fields of view, while those of mice were obtained at 40× magnification. Confocal images were processed with OlyVIA software (OLYMPUS OlyVIA3.3, Olympus, Japan). The mean fluorescence intensities of Nrf2 and NOX4 were quantified by automatic thresholding of the fluorescence intensity by using ImageJ software (NIH, USA). The number of Nrf2+/GFAP+ astrocytes and NOX4+/GFAP+ astrocytes were counted by an individual blinded to the sample identity. The percentages of Nrf2+/GFAP+ astrocytes and NOX4+/GFAP+ astrocytes among the total GFAP+ astrocytes were analysed. Thirty astrocytes were counted per human brain tissue section, whereas 80 astrocytes were counted per mouse brain tissue section. Morphometric analysis was performed on these astrocytes from both human and mouse brain slices. The number of branches/astrocytes and the average branch length/astrocytes were computed.

### 4-hydroxynonenal (4-HNE), 8-hydroxydesoxyguanosine (8-OHdG) and 3-nitrotyrosine (3-NT)

For immunofluorescence staining in mAS, cells grown on coverslips were washed with PBS and fixed with 4% paraformaldehyde for 20 min, followed by incubation for 10 min in 0.3% Triton X-100 (in PBS) to permeabilize the cells. The fixed mAS cells were incubated in blocking buffer (10% goat serum) for 60 min at room temperature, followed by incubation with primary antibodies, including anti-3-NT, anti-4-HNE, anti-8-OHdG, and anti-Tomm20 (**STable. 1**) at 4°C overnight. After three washes with PBS, the sections were incubated for 1 h at room temperature with AlexaFluor-546-conjugated donkey anti-rabbit IgG (1:500, Invitrogen). Sections were then mounted using vector anti-fading mounting medium. Fluorescence was detected by a confocal microscope (Olympus, Japan), and images were obtained at 100× magnification in randomly selected fields of view. Confocal images were processed with OlyVIA software (OLYMPUS OlyVIA3.3, Olympus, Japan).

The mean fluorescence intensities of 3-NT, 4-HNE, 8-OHdG and Tomm20 were quantified by automatic thresholding of the fluorescence intensity in these fluorescent images by using ImageJ software as described earlier [38, 39]. Mitochondrial morphology was used to analyse mAS cells. Mitochondria in normal mAS cells were filamentous and linear-tubular, whereas mitochondria in mitochondria-fragmented mAS cells were shortened, punctate and round. mAS cells with more than 70% mitochondrial fragmentation were identified as mitochondrial fragmented mAS cells. Each experiment was repeated three times, and at least 60 mAS cells per group were analysed.

### Morphological analysis of astrocytes

We analysed the morphology of astrocytes in both human and mouse brain tissue slices by using the method described by Young et al [40]. Briefly, representative immunofluorescence images with only GFAP-positive staining were analysed using ImageJ software. After adjusting the grayscale and contrast, removing background noise and binarizing the images, we obtained images of astrocytic processes. The skeleton map of astrocytes was then obtained using the plugin AnalyseSkeleton (2D/3D). We measured and calculated the number of branches and the average branch length for each cell. The data were averaged from at least 20 cells per group.

### Immunoblotting analysis

mAS cells were homogenized in ice-cold RIPA lysis buffer containing freshly added protease inhibitor (Sigma) [41]. Samples (40 μg protein/lane) were separated by 10% Bis-Tris gels and semidry transferred to polyvinylidene fluoride (PVDF) membranes (0.45 μm pore size, Millipore). After being blocked by incubation with TBST containing 5% nonfat milk, the membranes were incubated for 1 h at room temperature followed by overnight incubation at 4°C with primary antibodies, including anti-Nrf2, anti-HO-1, anti-GPX4, and anti-xCT (**STable. 1**). Glyceraldehyde-3-phosphate dehydrogenase (GAPDH) was used as an endogenous control. Subsequently, the membranes were washed three times with TBST (20% Tween-20 added in Tris buffered saline) and incubated with secondary antibody for 2 hours. An imaging system (SYNGENE, UK) was used to evaluate the protein expression, and the band density was analysed by ImageJ software.

### Detection of ROS

The level of ROS was detected following the manufacturer’s instructions [42]. Briefly, stably infected mAS cells were incubated with 5 μM dihydroethidium (DHE) for 30 min at 37°C. mAS cells were rotated gently and washed and resuspended in PBS. The mean fluorescence intensity was detected by a NovoCyte flow cytometer (Agilent Technologies, Santa Clara, USA). Data were analysed by NovoExpress software.

### Statistical analysis

All data are presented as the mean ± standard deviation (SD). All statistical analyses were performed using a two-tailed Student’s t test for comparison of two groups (GraphPad Prism version 8.0, GraphPad Software Inc., San Diego, CA, USA). A p value ≤0.05 was considered statistically significant.

## Results

### The expression levels of Nrf2 are decreased in astrocytes of the frontal cortex from patients with Alzheimer’s disease

First, we examined the expression levels of astroglia Nrf2 and the number of Nrf2/GFAP+ astrocytes in the frontal cortex from patients with AD and nondemented controls (n = 3 per group, **Figs. 1A-D**). The details of the demographics and pathological information of AD patients and nondemented controls are summarized in **Table 1**. Nrf2/GFAP double immunofluorescence staining was performed in frontal cortical slices.

**Figure 1.**
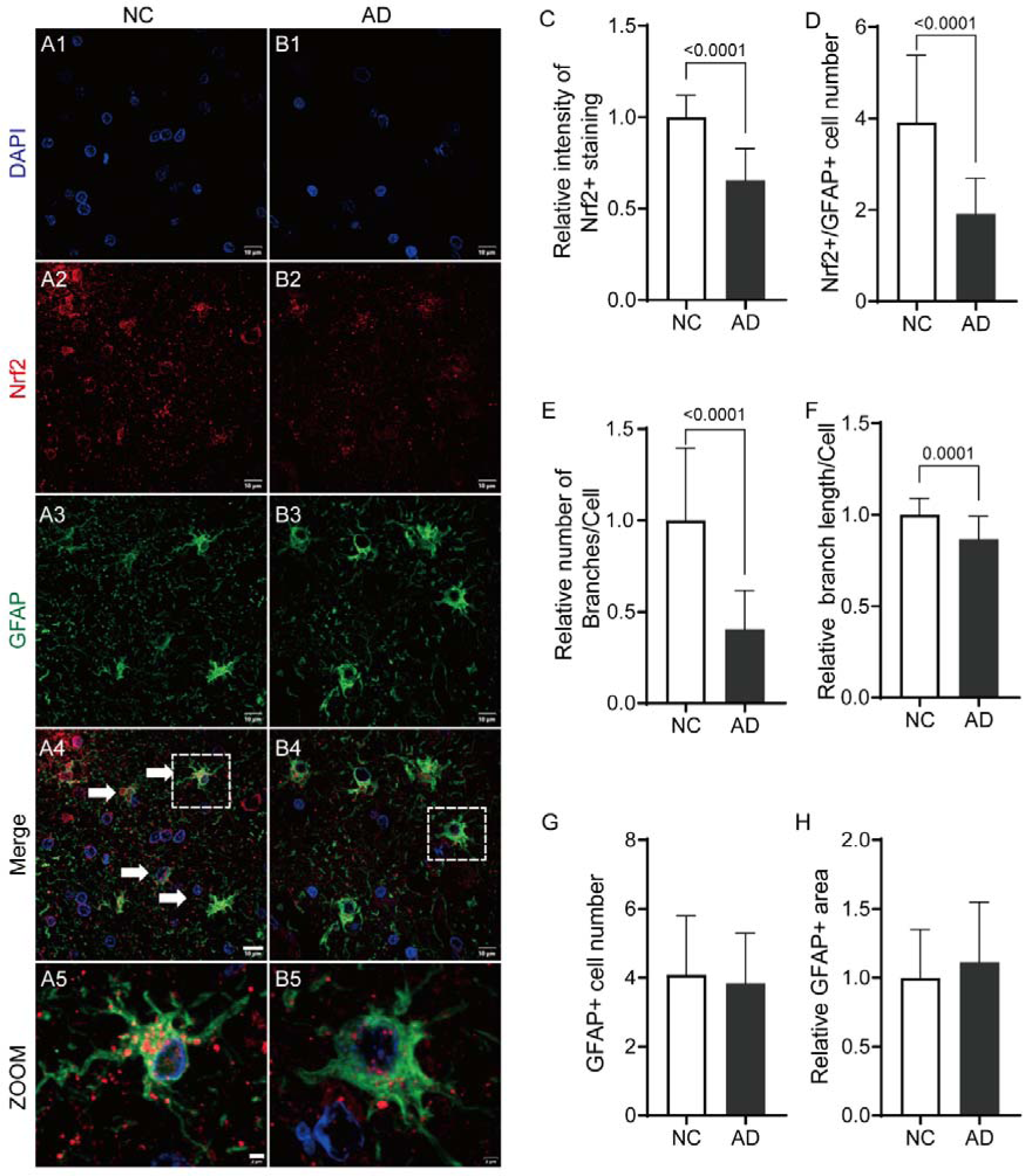
Decreased levels of astroglial Nrf2 in the frontal cortex from patients with AD. (A, B) Representative immunofluorescence images of Nrf2 (red) and GFAP (green) in the frontal cortex from patients with AD and nondemented controls (NCs). Nuclei are counterstained with DAPI (blue); Scale bar: 10 μm (row 1-4), 2 μm (row 5); Cells stained positive for Nrf2 and GFAP are indicated with white arrows. (C) Quantification of the fluorescence intensity of Nrf2 staining in astrocytes, (D) number of Nrf2+/GFAP+ astrocytes in the frontal cortex of patients with AD and NCs (n = 3 per group). (E) Reduced relative number of branches and (F) reduced branch length in astrocytes in the frontal cortex of patients with AD and NCs (n = 3 per group). (G) Number of GFAP+ astrocytes and (H) relative GFAP+ area in the frontal cortex of patients with AD and NCs (n = 3 per group). Data are expressed as the mean ± SD.

The intensity of Nrf2+ staining was decreased by approx. 40% in the frontal cortex of AD patients compared to nondemented controls (**Fig. 1C**, AD vs. NC; p<0.0001). The number of Nrf2+/GFAP+ astrocytes was decreased from 100% in the frontal cortex from nondemented controls to approx. half in AD patients (**Fig. 1D**, AD vs. NC; p<0.0001). GFAP+ astrocytes were shrunken in the frontal cortex from patients with AD compared to that from nondemented controls. The relative number of branches of GFAP+ astrocytes in the frontal cortex was reduced by 60% from AD compared to that from nondemented controls (**Fig. 1E**, AD vs. NC; p <0.0001). The relative branch length of GFAP+ astrocytes in the frontal cortex was reduced by approx. 14% in AD patients compared to that in nondemented controls (**Fig. 1F**, AD vs. NC; p = 0.0001). The number of GFAP+ astrocytes and GFAP+ area remained comparable in the frontal cortex from AD patients compared to that from nondemented controls (**Fig. 1G, H**).

### Increased NOX4 in astrocytes of frontal cortex from patients with Alzheimer’s disease

NADPH oxidases (NOXs) are a family of enzymes that produce ROS, and NOX4 is a major isoform in astrocytes [43]. To investigate the role of NOX4 in astrocytes from the brains of AD patients, we measured the levels of NOX4 expression in GFAP+ astrocytes and the number of NOX4+ astrocytes from the frontal cortex of AD patients and nondemented controls by immunofluorescence staining (**Figs. 2A-D)**. Different morphologies of astrocytes with fewer branches or only the soma were observed in the frontal cortex of AD brain tissue (**Figs. 2B-D**). We showed that the intensity of NOX4+ staining was elevated by approx. 50% in the frontal cortex of AD patients compared with nondemented controls (**Fig. 2E**, AD vs. NC, p < 0.0001). The number of NOX4+/GFAP+ astrocytes was approx. double in the frontal cortex of AD patients compared to nondemented controls (**Fig. 2F**, AD vs. NC, p < 0.0001). Increased subcellular colocalization of NOX4 with GFAP+ (96% astrocytes NOX4+) was observed in the frontal cortex of AD patients (**Figs. 2A-D**). Morphology analysis indicated a reduced relative number of branches and lower average branch length in astrocytes in the frontal cortex from AD patients compared to those in nondemented controls (**Figs. 2G, H**, AD vs. NC; p < 0.0001, p = 0.0305).

**Figure 2.**
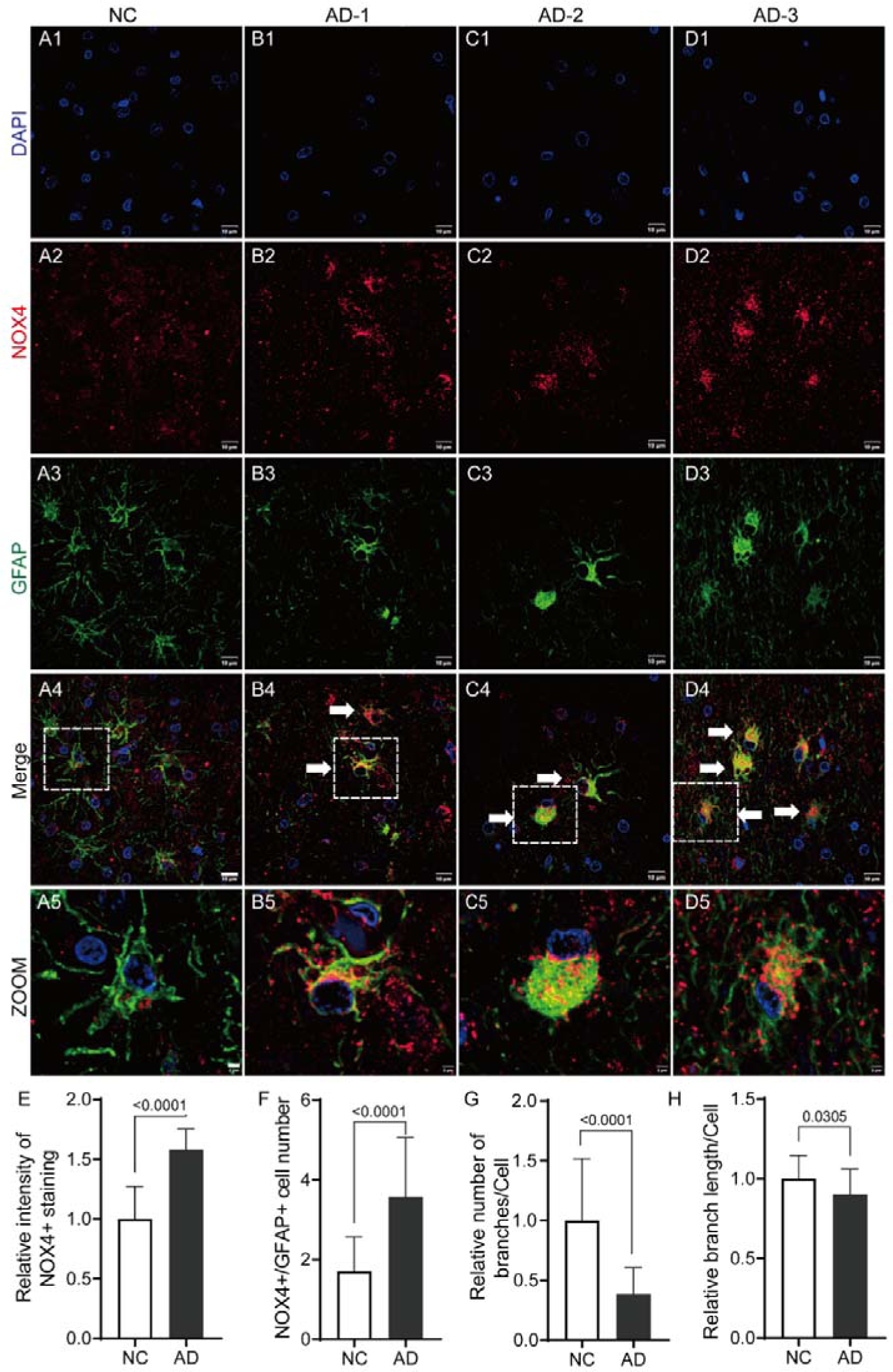
Increased levels of astroglial NOX4 in the frontal cortex from patients with AD. (A-B) Representative immunofluorescence images of NOX4 (red) and GFAP (green) in the frontal cortex from patients with AD (AD-1, AD-2, AD-3) and nondemented controls (NCs). Nuclei are counterstained with DAPI (blue); Scale bar: 10 μm (row 1-4), 2 μm (row 5); Cells stained positive for NOX4 and GFAP are indicated with white arrows. (C) Quantification of the fluorescence intensity of NOX4 staining in astrocytes, (D) number of Nrf2+/GFAP+ astrocytes, (E) reduced relative number of branches and (F) reduced branch length in astrocytes in the frontal cortex of patients with AD and NCs (n = 3 per group). (G) Comparable number of GFAP+ astrocytes in the frontal cortex of patients with AD and NC (n = 3 per group). Data are expressed as the mean ± SD.

### Nrf2 levels are decreased in astrocytes in the cortex of 3×Tg mouse brains

We further assessed the alterations of Nrf2 in the impairment of astrocytes in the 3×Tg mouse model of AD (at 10 months of age). The intensity of Nrf2 was decreased in GFAP+ astrocytes in the cortex from 3×Tg mice compared with age-matched wild-type (WT) mice using immunofluorescence staining (**Figs. 3A-C**, 3×Tg vs. WT, p = 0.0041). We found that the number of Nrf2+/GFAP+ astrocytes was reduced in the cortical region of 3×Tg mice compared with that of WT mice (**Figs. 3A, B, D**, 3×Tg vs. WT, p < 0.0001). The percentage of Nrf2+ astrocytes was reduced from 79% in WT mice to 41% in 3×Tg mice. The relative number of branches of GFAP+ astrocytes was reduced by approx. 20% in the cortical region of 3×Tg mice than in that of WT mice (**Fig. 3E**, 3×Tg vs. WT, p = 0.0429). The relative branch length of GFAP+ astrocytes was reduced by approx. 20% in the cortical region of 3×Tg mice than in that of WT mice (**Fig. 3F**, 3×Tg vs. WT, p = 0.0397).

**Figure 3.**
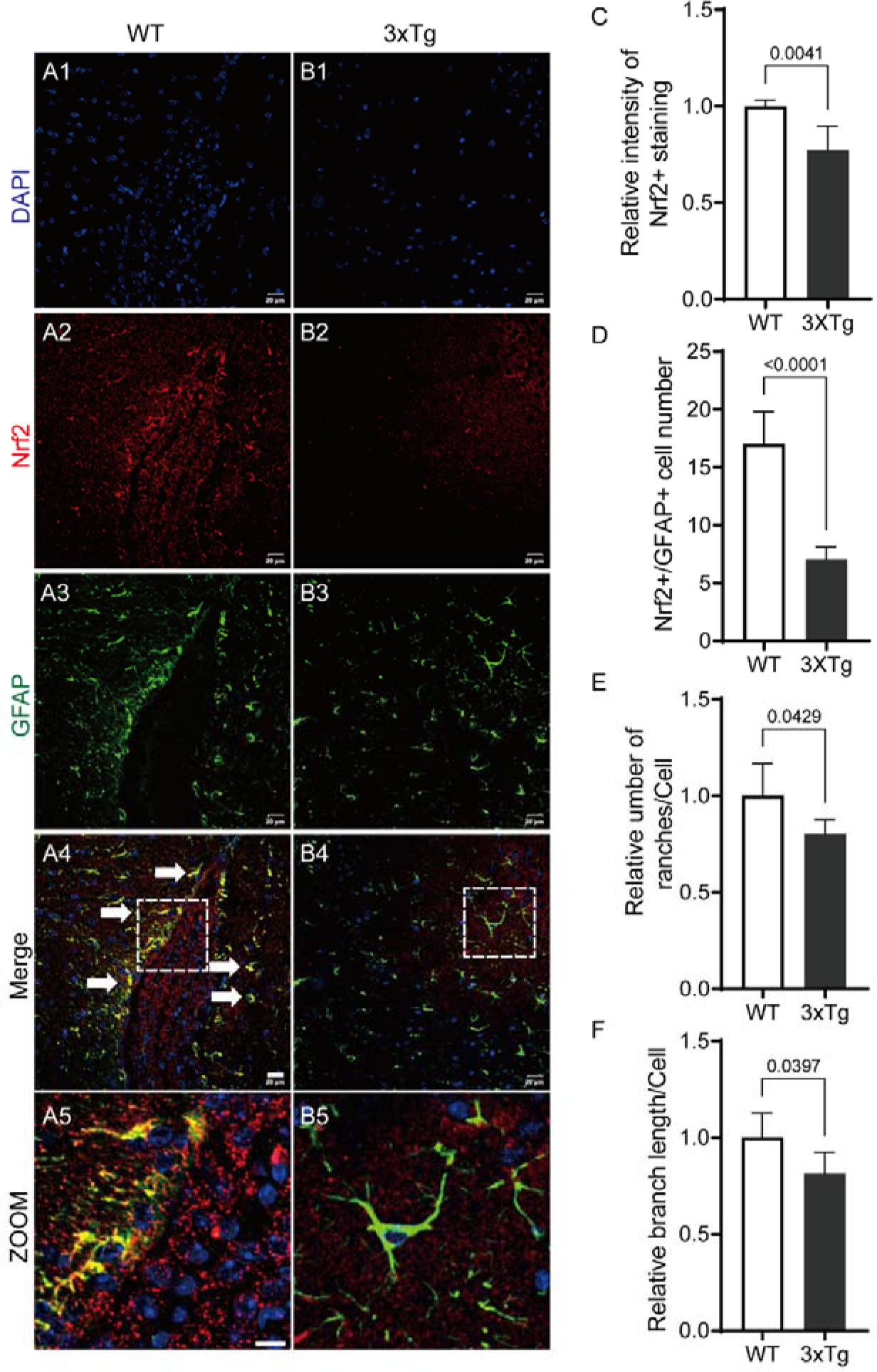
Decreased levels of astroglial Nrf2 in the cortex from 3×Tg mice. (A-B) Representative immunofluorescence images of Nrf2 (red) and GFAP (green) in the cortex from 3×Tg and wild-type (WT) mice. Nuclei are counterstained with DAPI (blue); Scale bar: 20 μm (row 1-4), 10 μm (row 5); Cells stained positive for Nrf2 and GFAP are indicated with white arrows. (C) Quantification of the fluorescence intensity of Nrf2 staining in astrocytes, (D) number of Nrf2+/GFAP+ astrocytes in the cortex from 3×Tg and WT mice (n = 5 per group). (E) Reduced relative number of branches and (F) branch length in astrocytes in the cortex from 3×Tg and WT mice (n = 5 per group). Data are expressed as the mean ± SD.

### The levels of NOX4 are increased in astrocytes of the cortex from 3×Tg mouse brains

Next, we analysed the protein levels of the ROS marker NOX4 in the cortex of 3×Tg mice (10 months old) compared to age-matched WT mice. We found that the intensity of NOX4 was increased in GFAP+ astrocytes in the cortex from 3×Tg mice compared to WT mice by immunofluorescence staining (**Figs. 4A-C**, 3×Tg vs. WT, p = 0.0079). Moreover, the number of GFAP+ astrocytes that colocalized with NOX4 was significantly increased in 3×Tg mice compared with WT mice (**Fig. 4D**, 3×Tg vs. WT, p = 0.0079). The percentage of NOX4+ astrocytes was 70% in the cortex of 3×Tg mice, which was higher than the 39% in WT mice. Morphology analysis indicated a reduced relative number of branches and lower average branch length in the astrocytes in the cortex from 3×Tg mice compared to WT mice (**Figs. 4E, F**, 3×Tg vs. WT, p = 0.0312, p = 0.0071). The total number of GFAP+ astrocytes in the cortex was comparable between 3×Tg mice and age-matched WT mice (**Fig. 4G**).

**Figure 4.**
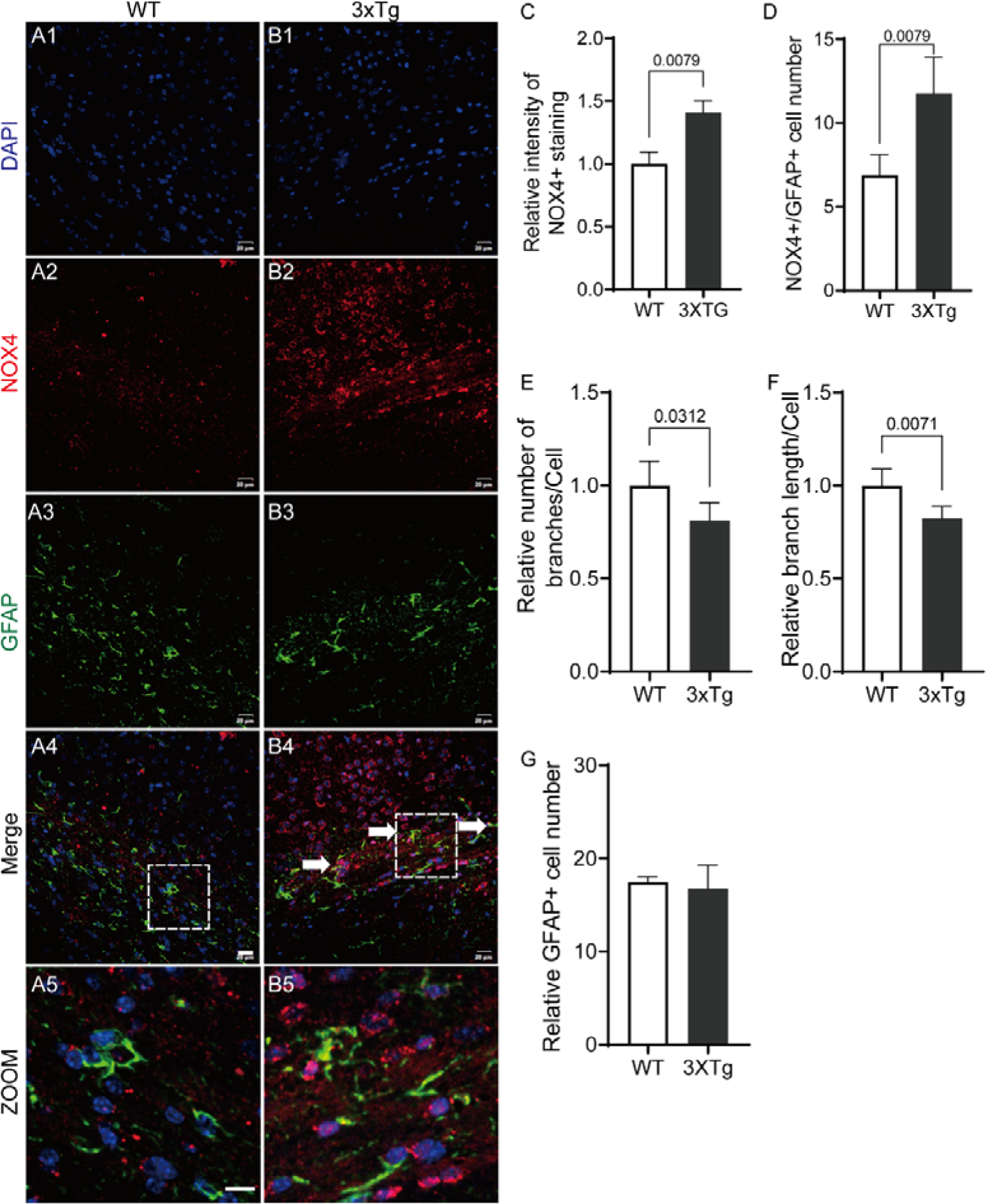
Increased levels of astroglial NOX4 in the cortex from 3×Tg mice. (A-B) Representative immunofluorescence images of NOX4 (red) and GFAP (green) in the cortex from 3×Tg and wild-type (WT) mice. Nuclei are counterstained with DAPI (blue); Scale bar: 20 μm (row 1-4), 10 μm (row 5); Cells stained positive for NOX4 and GFAP are indicated with white arrows. (C) Quantification of the fluorescence intensity of NOX4 staining in astrocytes, (D) number of NOX4+/GFAP+ astrocytes in the cortex from 3×Tg and WT mice (n = 5 per group). (E) Reduced relative number of branches and (F) reduced branch length in astrocytes in the cortex from 3×Tg and WT mice (n = 5 per group). Data are expressed as the mean ± SD.

### Nrf2 deficiency mediated HO-1/GPX4/xCT-related protein expression in mouse astrocytes

Next, we examined whether Nrf2 deficiency could affect redox homeostasis and the mechanisms of increased ferroptosis in mouse astrocytes. Ferroptosis is biochemically characterized by intracellular accumulation of iron and the accumulation of lipid peroxides, which can damage the lipid bilayer membrane through accelerated oxidation of membrane lipids [44]. Mouse astrocytes were infected with the Nrf2 shRNA-delivering lentivirus, with GFP fluorescence as the reporter. The GPX4/System Xc− pathway plays a key role in the downregulation of ferroptosis [45, 46]. GPX4 utilizes glutathione as a cofactor to repair oxidative damage to lipids and inhibits ferroptosis [47, 48]. System Xc− is composed of a light-chain subunit (xCT, SLC7A11) and a heavy-chain subunit (CD98hc, SLC3A2)[49]. We performed western blotting to measure the protein expression of HO-1/GPX4/xCT-related proteins in mouse astrocytes. We found that mouse astrocytes treated with shRNA showed decreased expression of Nrf2 by more than 60% compared with the control group (**Fig. 5B**, p = 0.0035). We also showed that the downregulation of Nrf2 inhibited the protein expression of HO-1 by 11% (**Fig. 5C**, p=0.0434) and GPX4 by 44% (**Fig. 5E**, p = 0.0028, vs. control) compared to the control. Moreover, the levels of xCT were increased by 20% in Nrf2-silenced mouse astrocytes compared to control astrocytes (**Fig. 5D**, p = 0.0408).

**Figure 5.**
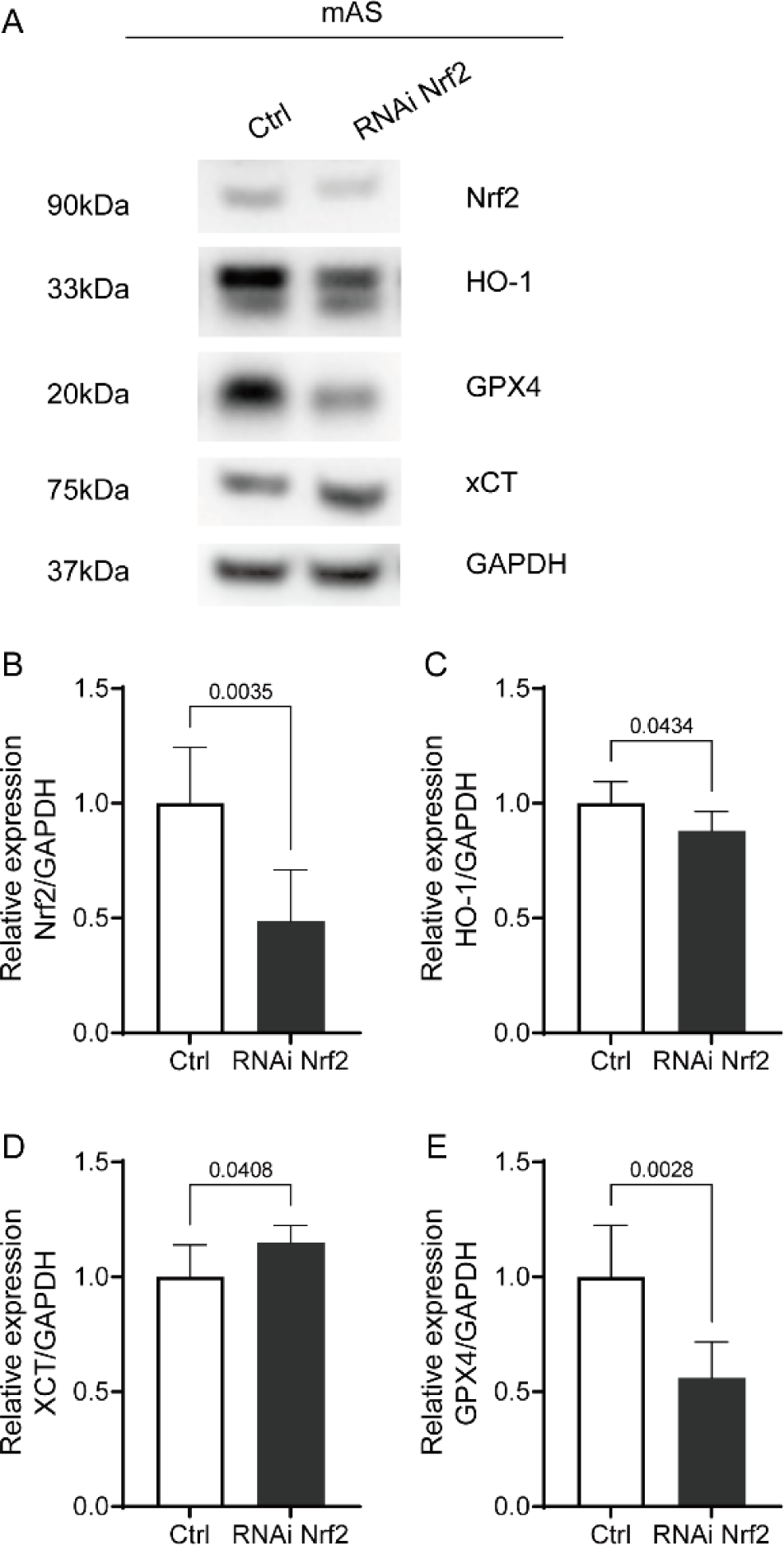
Nrf2 deficiency mediated HO-1/GPX4/xCT-related protein expression in mouse astrocytes. (A) Representative immunoblot images of Nrf2, HO-1, GPX4 and xCT in Nrf2-silenced mouse astrocytes and in the control group. (B-E) Quantification of the expression of Nrf2, HO-1, GPX4 and xCT in Nrf2-silenced mouse astrocytes and in the control group (n = 6 per group). Data are presented as the mean ± SD.

### Nrf2 Silencing promotes fragmentation of mitochondria and ferroptosis by oxidative stress in mouse astrocytes

Here, we analysed the fragmentation of mitochondria through immunofluorescence staining with the outer mitochondrial membrane 20 (TOMM20) (**Figs. 6A-C**). Immunofluorescence staining revealed that inhibition of Nrf2 induced the fragmentation of mitochondria; the percentage of the number of mAS cells with fragmentation of mitochondria was increased from 20% to 70% by Nrf2 silencing relative to the control (**Fig. 6C,** p = 0.0019, vs. control).

**Figure 6.**
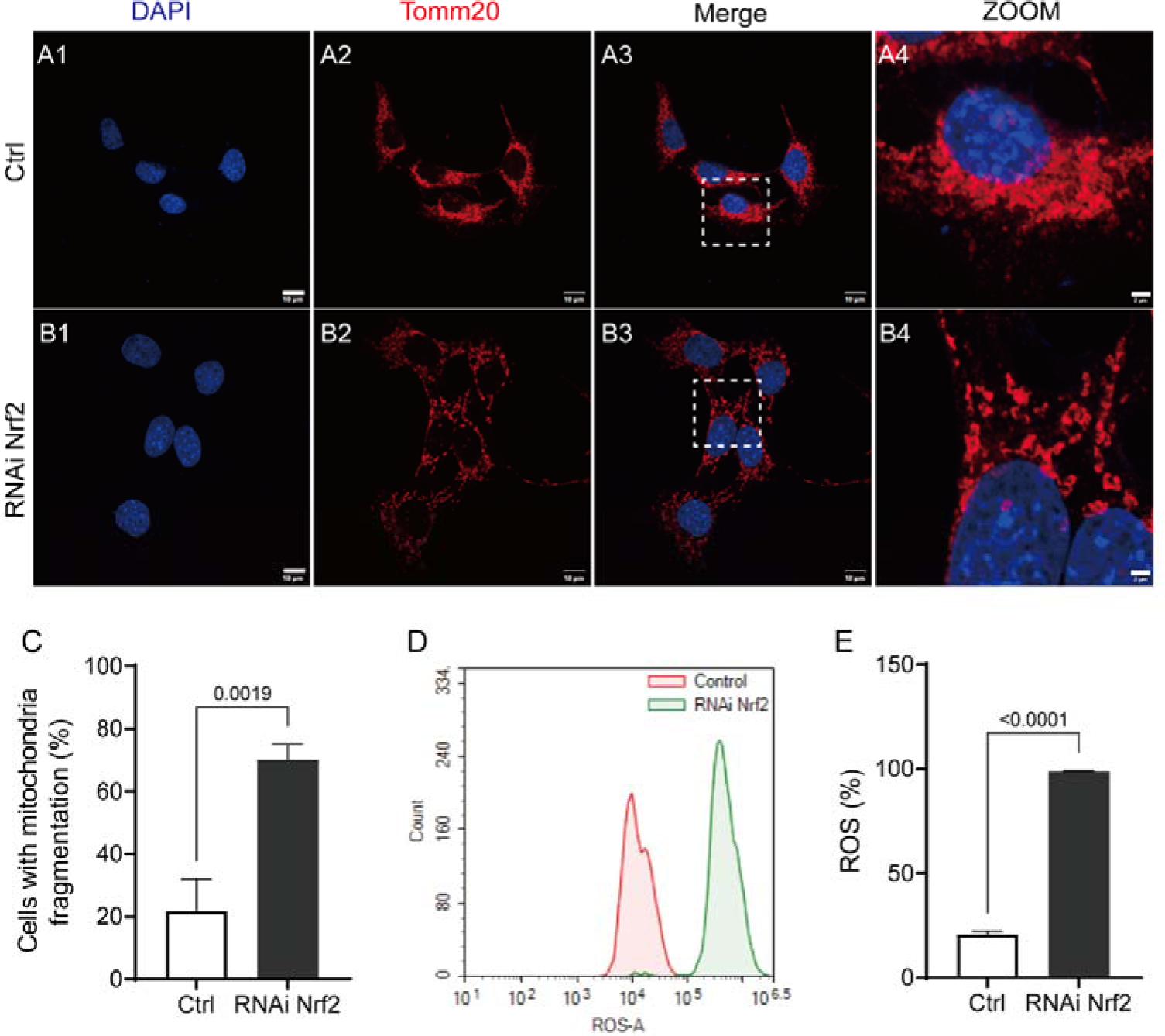
Nrf2 deficiency promoted ferroptosis by oxidative stress in mouse astrocytes. (A, B) Representative confocal images of Tomm20 immunofluorescence (red) in Nrf2-silenced mouse astrocytes in the control group. Nuclei were counterstained with DAPI (blue); Scale bar: 10 μm. 2 μm (zoom). (C) Quantitation of the fluorescence intensity of Tomm20 staining in Nrf2-silenced mouse astrocytes in the control group (n = 3 per group). (D, E) Increased ROS levels in Nrf2-silenced mouse astrocytes compared to the control by flow cytometry using the DHE assay (n = 3 per group). Data are expressed as the mean ± SD.

To investigate the potential role of Nrf2 in oxidative stress in mouse astrocytes, we examined alterations in ROS after Nrf2 silencing. A fluorescence DHE flow cytometry assay was used to measure the production of ROS in mouse astrocytes (**Fig. 6D**). DHE signal intensity, which is indicative of ROS levels, was 80% higher in Nrf2-silenced mouse astrocytes than in control mAS cells (**Fig. 6E**, p<0.0001).

Next, we examined alterations in lipid peroxidation and DNA oxidation after Nrf2 silencing. To measure oxidative damage, products of oxidative stress, such as 4-HNE from lipid peroxidation and 8-OHdG from DNA oxidation, were measured using immunofluorescence staining in mouse astrocytes after Nrf2 silencing (**Fig. 7**). 4-HNE, an electrophilic lipid peroxidation product, has various cytotoxic effects on lipid peroxidation, such as depletion of glutathione and reduction in enzyme activities [50]. The fluorescence intensity of 4-HNE (**Figs. 7A-C**, p = 0.0076) and 8-OHdG (**Figs. 7D-F**, p = 0.0267) was increased in Nrf2-silenced mouse astrocytes compared to the control group. Furthermore, we measured 3-NT in mouse astrocytes, which is an indicator of posttranslational oxidative damage. We showed significantly elevated 3-NT expression in Nrf2-silenced mouse astrocytes compared to control astrocytes (**Figs. 7G-I**, p = 0.043), supporting that Nrf2 deficiency increased tyrosine nitration in mouse astrocytes.

**Figure 7.**
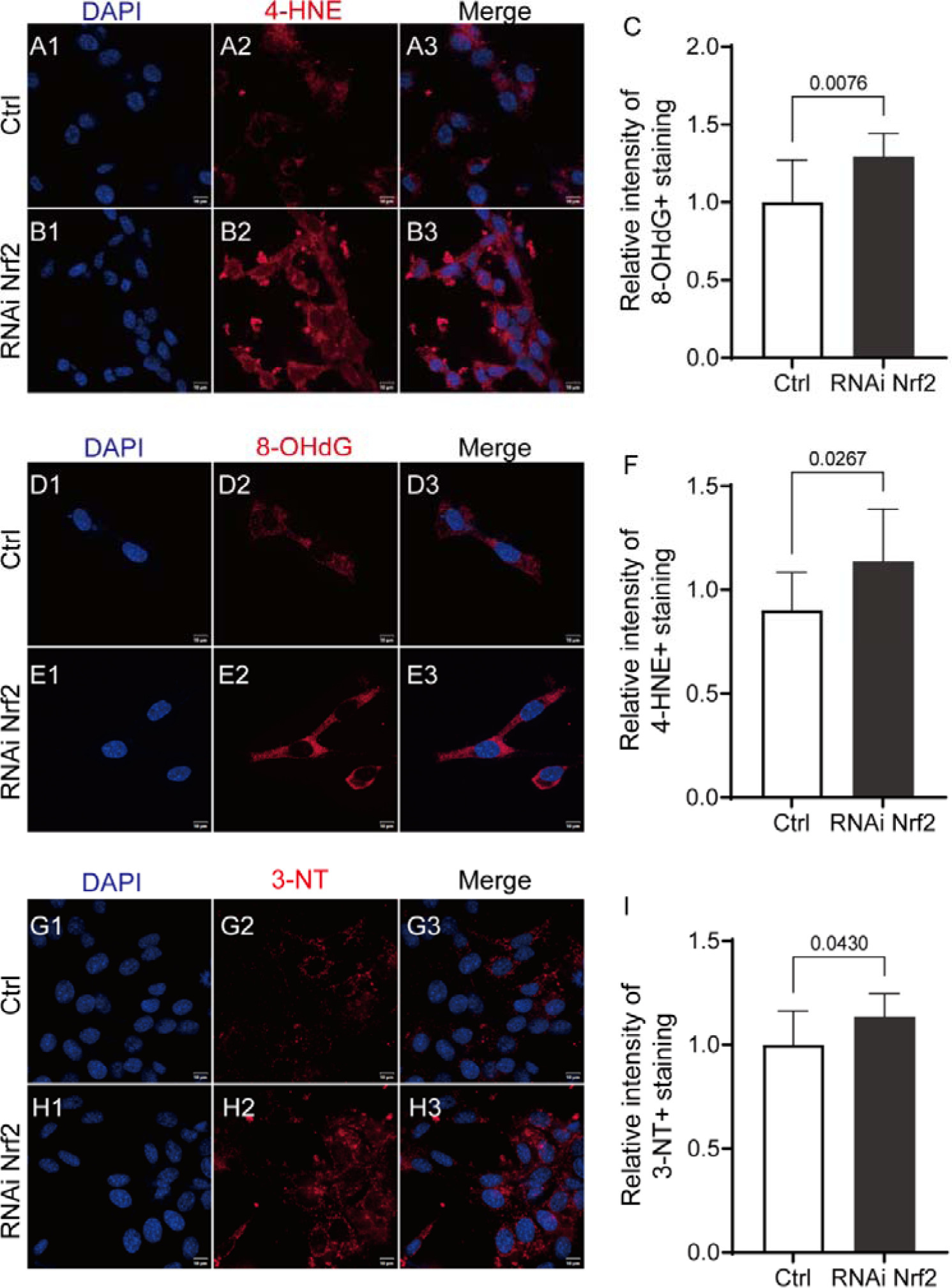
Nrf2 deficiency promoted ferroptosis by oxidative stress in mouse astrocytes. (A, B) Representative confocal images of the immunofluorescence of 4-HNE (red) in Nrf2-silenced mouse astrocytes in the control group. Nuclei were counterstained with DAPI (blue); Scale bar: 10 μm. (C) Quantitation of the fluorescence intensity of 4-HNE staining in Nrf2-silenced mouse astrocytes in the control group (n = 3 per group). (D, E) Representative confocal images of the immunofluorescence of 8-OHdG (red) in Nrf2-silenced mouse astrocytes in the control group. Nuclei were counterstained with DAPI (blue); Scale bar: 10 μm. (F) Quantitation of the fluorescence intensity of 8-OHdG staining in Nrf2-silenced mouse astrocytes in the control group (n = 3 per group). (A, B) Representative confocal images of the immunofluorescence of 3-NT (red) in Nrf2-silenced mouse astrocytes in the control group. Nuclei were counterstained with DAPI (blue); Scale bar: 10 μm. (C) Quantitation of the fluorescence intensity of 3-NT staining in Nrf2-silenced mouse astrocytes in the control group (n = 3 per group). Data are expressed as the mean ± SD.

## Discussion

In the current study, we demonstrated a decreased level of Nrf2 and an elevated level of NOX4 in astrocytes in patients with AD and a 3×Tg mouse model of AD. We found that Nrf2 deficiency induced ROS production, mitochondrial impairment, ferroptosis-dependent cytotoxicity involves the activation of lipid peroxidation and DNA oxidation in mouse astrocytes.

Increasing evidence suggests that excessive ROS levels [51] and oxidative stress play important roles in the pathogenesis of AD and various AD animal models, including 3×Tg mice [52-54]. Oxidative stress may occur as a result of increased free radicals and decreased antioxidant defences [55, 56]. The brain is particularly vulnerable to free radical damage compared with other organs [57]. Free radicals are necessary for synaptic plasticity and learning and memory under physiological conditions; on the other hand, increased oxidative stress impairs synaptic plasticity and memory in AD [58, 59]. Excessive ROS production can result in the accumulation of Aβ plaques and hyperphosphorylated tau, which are the main pathological hallmarks of AD [10].

Here, we observed a reduction in the level of Nrf2 and an elevation in the level of NOX4 in astrocytes in the frontal cortex of AD patients, in the cortex of 3×Tg mice, and in mouse astrocytes. This finding is in line with previous reports of Nrf2 reduction in cortical brain tissue from AD patients [24] and in brain tissue from 3×Tg mice [60]. Elevation of the level of NOX4 and its involvement in promoting the ferroptosis of astrocytes has been reported in the postmortem brain from AD and APP/PS1 mice [24]. Attenuation of Nox4 by using Nox4 targeting miR-204-3p and the inhibitor GLX351322 protected neuronal cells against Aβ-induced neurotoxicity, decreased oxidative stress and alleviated memory deficits in APP/PS1 mice, although GFAP was not changed after Lv-miR-204 treatment [61]. It is noted that in addition to astrocytes, the effect of Nrf2 activation has been shown to protect neuronal cells from Aβ-mediated oxidative and metabolic damage mediated through the PI3K/GSK-3 axis [33] as well as via BACE-1 pathway [34]. Previous studies have reported complex roles for reactive astrocytes [62, 63], affecting hyperphosphorylation and aggregation of tau and Aβ pathology clearance [3, 64]. In the current study we focused on the astrocyte involvement in ferroptosis and oxidative stress in AD.

We showed that Nrf2 deficiency increased the levels of xCT and decreased the levels of GPX4 in mouse astrocytes. Impaired GPX4/xCT perturbs glutathione synthesis and cellular redox balance and induces ferroptosis [45, 46]. Moreover, System Xc− takes up cystine into the cell in exchange for glutamate [49]. Inhibition of System Xc− reduces glutathione levels and impairs GPX4 activity, thereby increasing lipid peroxidation [65]. While it transports intracellular glutamate to the extracellular space, System Xc− transports extracellular cysteine into the cell [66]. The inhibition of System Xc− thus leads to a compensatory transcriptional upregulation of xCT expression in cells [67].

We found that the decrease in Nrf2 expression in astrocytes in AD was accompanied by elevated levels of oxidative stress in mouse astrocytes. Moreover, Nrf2 silencing in mouse astrocytes increased ROS generation (DHE), lipid peroxidation (4-HNE), DNA oxidation (8-OHdG), and posttranslational oxidative damage (3-NT). ROS and free lipid radicals are generated when anti-ferroptosis defence pathways are interrupted in cells [68-70]. Under ferroptotic stress, ROS production and lipid peroxidation generate toxic products that damage nucleic acids and cellular proteins, resulting in cell death [71, 72]Lipid peroxidation induced by ROS represents the state of oxidative stress, which could trigger ferroptosis [73]. High levels of 4-HNE are involved in the oxidative stress of AD or Parkinson’s disease (PD), associated with cellular stress and pro-apoptosis [24]. Previous studies reported increased levels of 8-OHdG in mitochondrial brain fractions in AD brain compared with that from nondemented controls [74, 75]. Excessive levels of ROS can lead to the formation of posttranslational protein tyrosine oxidation to yield 3-NT, which is a biologically relevant protein related to neurodegenerative processes [76]. Nrf2 could thus be an upstream molecular target in the impairment of oxygen metabolism in astrocytes in AD. In addition, Earlier studies showed that pharmacological upregulation of Nrf2 ameliorated ROS and cognitive deficits in 3×Tg mice [34]. Nrf2 deficiency increased Aβ plaques and tau accumulation, neuroinflammation, oxidative stress and cognitive impairment in APP/PS1 transgenic mice [77, 78]. Another study in an App knock-in AD mouse model showed that Nrf2 is linked to excessive ROS production and oxidative stress [79].

We found that Nrf2 silencing in mouse astrocytes increased mitochondrial fragmentation (Tomm20) in mouse astrocyte. Mitochondria play a critical role in the regulation of ferroptosis [80]. The integrity of the mitochondrial membrane is important for maintaining mitochondrial function and is impaired in AD [81-83]. Mitochondria supports neuronal activity by providing energy supply, as well as protects neurons by minimizing mitochondrial related oxidative damage [84]. Malfunction in the regulation of astrocytic mitochondrial dynamics may impair the capability of mitochondria to regulate metabolic demand, and disturb glial-neuronal interactions [85]. Microglial Mitochondrial fragmentation has been shown to trigger the A1 astrocytic response and lead to inflammatory neurodegeneration [86, 87]. Recent study showed the association between alterations of bioenergetics, mitochondria-ER interactions and proteostasis in hippocampal astrocytes from 3×Tg mice [88].

There are several limitations in our study. First, we investigated the cortex of both human and mouse brains. Given the regional diversity of astrocytes, further analysis of the mechanism underlying ferroptosis in astrocytes in the hippocampus is warranted. Second, further comprehensive approaches, such as transcriptomic studies, are needed to better profile the heterogeneity and distinctive molecular signatures of astrocytes [5]. Moreover, only male 3xTg mice were investigated in the current study. Given the impact of sex and hormone on oxidative stress and AD; further analysis in the female mouse models of AD are needed [89].

## Conclusion

In conclusion, our study demonstrates that decreasing Nrf2 promotes ferroptosis of astrocytes involving oxidative stress-induced ROS, mitochondrial impairment and lipid peroxidation as crucial mechanisms in AD. Our findings support the need for further study of Nrf2 as a therapeutic target for AD.

## Author Contributions

ZT, RN, and XLQ contributed to the conception and study design. ZYC and TZ performed the experiments. ZT, ZYC and YX contributed to data collection and data analysis. RN and ZT interpreted the data. XLQ, ZYC, RN, and ZT wrote the manuscript. All authors approved the manuscript before submission.

