## Supplemental Tables for "NRF2 deficiency promotes ferroptosis of astrocytes mediated by oxidative stress in Alzheimer’s disease"

**STable 1 Antibodies used in this study.**

| Antibody | Host | WB Dilution | IF Dilution | Sources | Catalog No. |
| --- | --- | --- | --- | --- | --- |
| Anti 3-NT | m |  | 1/50 | Santa-Cruz | #sc-32731 |
| Anti 4-HNE | r |  | 1/200 | Abcam | #ab46545 |
| Anti 8-OHdG | m |  | 1/200 | Abcam | #ab62623 |
| Anti GPX4 | r | 1/5000 |  | Abcam | #ab125066 |
| Anti xCT | m | 1/5000 |  | Abcam | #ab175186 |
| Anti GFAP | m |  | 1/200 | Cell Signaling Technology | #3670 |
| Anti Tomm20 | r |  | 1/300 | Abcam | #ab186734 |
| Anti GAPDH | r | 1/6000 |  | Genetex | #GTX100118 |
| Anti HO-1 | r | 1/1500 |  | Genetex | #GTX101147 |
| Anti NOX4 | r |  | 1/50 | Proteintech | #14347-1-AP |
| Anti Nrf2 | m | 1/4000 |  | Proteintech | #66504-1-Ig |
| Anti Nrf2 | r |  | 1/200 | Proteintech | #16396-1-AP |
| Donkey anti-Mouse IgG (H+L) Alexa Fluor™ 488 | d |  | 1/200 | Thermo Fisher Scientific | #A21202 |
| Donkey anti-Rabbit IgG (H+L) Alexa Fluor™546 | d |  | 1/200 | Thermo Fisher Scientific | #A10040 |
| Goat anti-Mouse IgG (H+L) Cyanine3 | g |  | 1/200 | Thermo Fisher Scientific | #A10521 |
| Goat anti-Mouse IgG (H+L) Secondary Antibody, HRP | g | 1/5000 |  | Thermo Fisher Scientific | #31430 |
| Goat anti-Rabbit IgG (H+L) Secondary Antibody, HRP | g | 1/5000 |  | Thermo Fisher Scientific | #31460 |

3-NT, 3-nitrotyrosine; 4-HNE, 4-hydroxynonenal; 8-OHdG, 8-hydroxy-2’-deoxyguanosine; m, mouse; d, donkey; g, goat; GAPDH, glyceraldehyde 3-phosphate dehydrogenase; GFAP, glial fibrillary acidic protein; GPX4, glutathione peroxidase 4; HO-1, Heme Oxygenase-1; HRP, horseradish peroxidase; IF, immunofluorescence; NOX4, nicotinamide adenine dinucleotide phosphate oxidase 4; Nrf2, nuclear factor E2 related factor 2; r, rabbit; WB, western blot; xCT, cystine/glutamate antiporter SLC7A11;

**STable 2 Reagents used in this study.**

| Regents | Sources | Catalog No. |
| --- | --- | --- |
| BIS-Tris gels | absin | #abs9391 |
| Fetal Bovine Serum | Biological Industries | #C2820-0500HI |
| Nonfat Dried Milk | Coolaber | #CN7861 |
| Dulbecco's Modified Eagle Medium | Gibco | #11965092 |
| Penicillin-streptomycin Solution | HyClone | #SV30010 |
| Hydrophobic PVDF Transfer Membrane | Merck Millipore Billerica | #ISEQ00010 |
| Immobilon Western HRP Chemiluminescent Substrate | Merck Millipore Billerica | #WBKLS0500 |
| TBS | Servicebio | #G0001 |
| Goat Serum | Solarbio | #SL038 |
| PMSF | Solarbio | #P0100 |
| Puromycin | Solarbio | #P8230 |
| RIPA buffer (high) | Solarbio | #R0010 |
| ROS assay kit | Solarbio | #CA1420 |
| Triton X-100 | Solarbio | #T8200 |
| Tween-20 | Solarbio | #T8220 |
| DAPI | SouthernBiotech | #0100-20 |
| Immunohistochemical Antigen Repair Buffer | ZSGB-BIO | #ZLI-9065 |

PVDF, polyvinylidene fluoride; TBS, tris buffered saline; PMSF, phenylmethylsulfonyl fluoride; RIPA, radio immunoprecipitation assay; ROS, reactive oxygen species; DAPI, 4′,6-diamidino-2-phenylindole.
